# Parallel processing by distinct classes of principal neurons in the olfactory cortex

**DOI:** 10.1101/2021.07.13.452206

**Authors:** Shivathmihai Nagappan, Kevin M. Franks

## Abstract

Understanding the specific roles that different neuron types play within a neural circuit is essential for understanding what that circuit does and how it does it. Primary olfactory (piriform, PCx) cortex contains two main types of principal neurons: semilunar (SL) and pyramidal (PYR). SLs and PYRs have different morphologies, connectivity, biophysical properties, and downstream projections, predicting distinct roles in cortical odor processing. The prevailing model is that odor processing in PCx occurs in two stages, where SLs are the primary recipients of olfactory bulb (OB) input, and PYRs receive and transform information from SLs. Here we recorded from optogenetically-identified SLs and PYRs in awake, head-fixed mice. We found differences in SL and PYR odor-evoked activity that reflect their different connectivity profiles. But SL responses did not precede PYR responses and suppressing SL activity had little effect on PYR odor responses. These results suggest that SLs and PYRs form parallel odor processing channels.

---

The brain contains a diverse array of neuron types that are each thought to play a specific role in neural circuit computations (Kim et al., 2020; Sanes and Masland, 2015; Yao et al., 2021). Understanding the specific role each neuron type plays will therefore be crucial to gaining a mechanistic understanding of what a neural circuit does and how it does it (Zeng and Sanes, 2017). PCx is a three-layered allocortex with principal neurons primarily located in layer II. There are two main types of layer II principal neurons: SLs in more superficial layer II and PYRs in deeper layer II. SLs and PYRs have distinct morphologies, input sources, intrinsic biophysical properties, and downstream projection targets (Diodato et al., 2016; Haberly, 1983; Heimer and Kalil, 1978; Mazo et al., 2017; Suzuki and Bekkers, 2006, 2011). These differences have motivated various predictions about the roles of SLs and PYRs in cortical odor processing, and how odor information is routed through PCx.

PCx is a sensory-associative cortex, containing both afferent OB inputs and recurrent or intracortical excitatory inputs (Franks et al., 2011; Hagiwara et al., 2012; Johnson et al., 2000; Suzuki and Bekkers, 2011; Wiegand et al., 2011). Mitral cells, which relay odor information from OB to PCx, send diffuse and overlapping projections along the lateral olfactory tract (LOT) into PCx (Buonviso et al., 1991; Ghosh et al., 2011; Miyamichi et al., 2011; Sosulski et al., 2011), allowing individual PCx neurons to sample information from multiple, unique combinations of glomeruli (Apicella et al., 2010; Davison and Ehlers, 2011). This provides PCx neurons the substrate to integrate elemental sensory information from the OB to form odor-specific neuronal ensembles and transform these ensembles through the recurrent, associative network (Barkai et al., 1994; Bolding and Franks, 2018; Bolding et al., 2020; Haberly, 2001; Pashkovski et al., 2020; Poo and Isaacson, 2011).

PYRs receive OB inputs on their distal apical dendrites, dense excitatory intracortical inputs on their proximal apical dendrites, and ‘top-down’ inputs from other brain regions on their basal dendrites. By contrast, SLs receive strong OB inputs on their distal apical dendrites, few if any excitatory intracortical inputs, and lack basal dendrites (Figure 1a). These differences in afferent and associative connections onto SLs and PYRs perhaps represent the two ends of the PCx sensory-associative continuum, where SLs simply combine afferent OB inputs while PYRs integrate and transform OB inputs through their intracortical connections. However, whether and how SLs and PYRs differentially encode odors is unknown.

**Figure 1.**
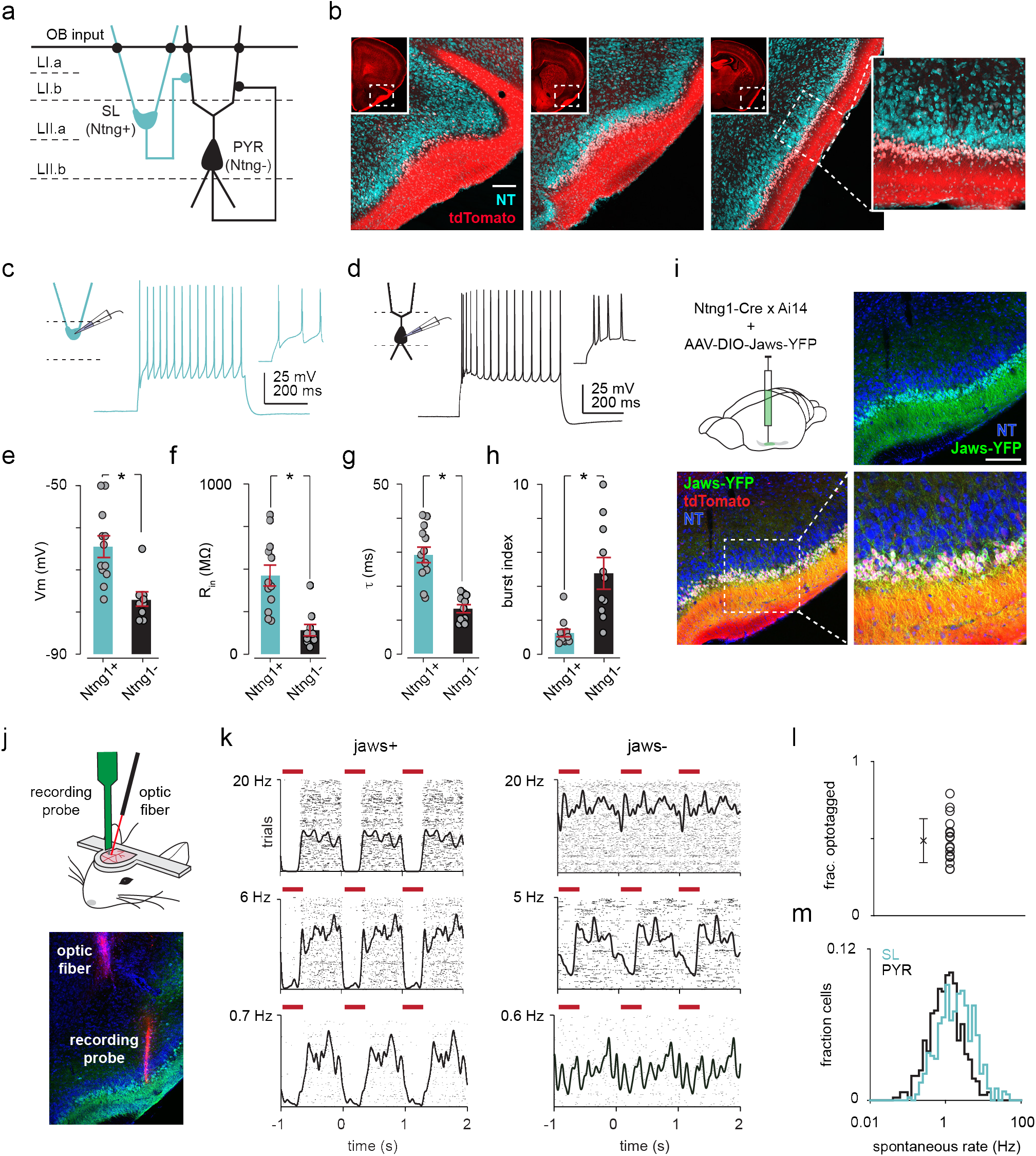
SLs can be optogenetically identified from extracellular recordings of awake, head-fixed Ntng1-Cre mice. (**a**) Schematic of SL and PYR connectivity and input sources within PCx. (**b**) Coronal sections from a Ntng1/Ai14 mouse brain showing tdTomato labeling from anterior to posterior PCx. tdTomato labeling is restricted to superficial Layer II of PCx. Scale bar is 200 μm. (**c**) Whole-cell current clamp recordings were obtained from acute brain slices isolated from Ntng1-Cre/Ai14 mice. Example voltage responses to direct current injections for a Ntng1+ (tdTomato+) cell. (**d**) As in (**a**) but for a Ntng1- (unlabeled) cell. (**e**) Resting membrane potentials of Ntng1+ and Ntng1- cells (Ntng1+: −65.4 ± 2.61 mV, n = 11 cells; Ntng1-: −77.0 ± 2.21 mV, n = 7 cells; p = 5.27 x 10^-4^, unpaired t-test). (**f**) Input resistances of Ntng1+ and Ntng1- cells (Ntng1+: 541 ± 68.8 MΩ, n = 9 cells; Ntng1-, 166 ± 38.9 MΩ, n = 8 cells; p = 2.15 x 10^-4^, unpaired t-test). (**g**) Membrane time constants of Ntng1+ and Ntng1- cells (Ntng1+: 32.1 ± 2.15 ms, n = 11 cells; Ntng1-: 13.6 ± 1.34 ms, n = 8 cells; p = 1.29 x 10^-5^, unpaired t-test). (**h**) Burst indices of Ntng1+ and Ntng1- cells (Ntng1+: 1.16 ± 0.130, n = 8 cells; Ntng1-: 4.57 ± 0.829, n = 8 cells; p = 1.43 x 10^-4^, unpaired t-test). (**i**) Coronal section of anterior PCx from an example Ntng1-Cre mouse showing selective and robust expression of the inhibitory opsin, Jaws, in SLs. Scale bar is 200 μm. (**j**) Schematic of probe and optic fiber positioning for opto-tagging (top). Histology showing positioning of the recording probe and optic fiber in PCx after both were painted with DiI (bottom). (**k**) Raster plots with trials aligned to the onset of each light pulse for six example units that were later categorized as either Jaws+ (left) or Jaws- (right). (**l**) Fraction of recorded cells from each experiment categorized as SLs (n=15 experiments). (**m**) Distribution of spontaneous spike rates for SLs (blue, n=426 cells) and PYRs (black, n=464 cells).

At one extreme, SLs and PYRs may form parallel and largely independent odor processing streams such as those that have been described in the auditory (Williamson and Polley, 2019), visual (Glickfeld et al., 2013), motor (Economo et al., 2018) and somatosensory cortices (Chen et al., 2015), where principal neurons projecting to distinct downstream regions encode different features of a stimulus or are differentially modulated during task learning. SLs and PYRs project to distinct downstream regions, lending support to a parallel processing model (Diodato et al., 2016; Mazo et al., 2017). Alternatively, odor processing in PCx may require the sequential activation of SLs and PYRs (Ketchum and Haberly, 1993; Suzuki and Bekkers, 2006, 2011; Wiegand et al., 2011), where SLs are the primary recipients of OB input, and their output is required to recruit PYRs. This sequential model is based on the observations that SLs receive stronger LOT inputs than PYRs, but little or no recurrent or intracortical inputs (Suzuki and Bekkers, 2006, 2011), and that SLs respond 2-4 ms earlier than PYRs following electrical stimulation of the LOT (Ketchum and Haberly, 1993; Suzuki and Bekkers, 2006, 2011; Wiegand et al., 2011). Finally, SLs and PYRs may be embedded within a parallel loop where mitral cells project onto both SLs and PYRs, and SLs additionally project onto PYRs, similar in organization to the entorhinal cortex (EC)-dentate gyrus (DG)-CA3 circuit in the hippocampus. Both DG and CA3 receive inputs from the EC. EC inputs into CA3 trigger the retrieval of information stored within the CA3 associative network while DG inputs into CA3 augment small differences in EC inputs to avoid similar EC inputs from recalling the same stored pattern (Leutgeb et al., 2007). Importantly, CA3 function is not impaired in the absence of DG inputs (Lee and Kesner, 2004).

In this study, we recorded extracellularly from layer II PCx neurons in awake, head-fixed Ntng1-Cre transgenic mice, which allowed us to distinguish SL activity from PYR. We first asked if and how the odor response properties of SLs and PYRs were different. If the dense intracortical inputs received by PYRs facilitate the stabilization of odor representations, then PYRs should be more odor selective, their responses less variable across trials and more discriminable between different odors. Second, we asked if odor information from OB is processed sequentially in PCx, in which case SL odor responses should precede PYR odor responses, and PYRs would require SL input for their activation. We found that SLs were more broadly tuned and odors were slightly more discriminable in PYRs than in SLs. SLs and PYRs responded with similar latencies and crucially, optogenetic and chemogenetic suppression of SLs had little effect on PYR odor responses. Together, these findings support a parallel processing model, where SLs and PYRs both receive direct OB inputs and relay different transformations of this sensory information to distinct downstream regions.

## RESULTS

### Ntng1-Cre transgenic mouse enables selective targeting of SLs *in vivo*

We generated a knock-in mouse line that expresses Cre-recombinase in Ntng1-expressing (Ntng1+) cells throughout the brain (Bolding et al., 2020) (Figure S1). In PCx, Cre-expressing cells were largely restricted to the superficial part of layer II (i.e. layer IIa), where SLs are located (Figure 1b). To confirm that the Cre-expressing cells were SLs, we sparsely labeled subsets of Ntng1+ cells by injecting a low-titer Cre-dependent adeno-associated virus (AAV) carrying green fluorescent protein (GFP) (Figure S2a). The large majority of GFP+ cells exhibited hallmark morphological characteristics of SLs: half-moon shaped somata, two prominent apical dendrites emerging directly from the cell body and no basal dendrites (fraction of morphologically identified SLs, mean ± s.d.: 87.9 ± 3.54% of labeled cells, n = 3 mice; Figure S2b). Additionally, we determined whether Ntng1+ cells exhibited the intrinsic biophysical properties described in SLs. We obtained whole-cell current-clamp recordings in acute brain slices isolated from Ntng1-Cre/Ai14 mice and compared the responses of tdTomato+ (i.e. Ntng1+) cells in layer IIa and unlabeled cells in layer IIb to direct current injections (Figures 1, c and d). We found that Ntng1+ and Ntng1- cells had different resting membrane potentials (Figure 1e), input resistances (Figure 1f), membrane time constants (Figure 1g), and burst indices (Figure 1, c, d and h), consistent with values reported for SLs and PYRs (Suzuki and Bekkers, 2006, 2011).

Previous studies have also reported that SLs receive few intracortical excitatory inputs and form excitatory synaptic connections onto PYRs and local inhibitory interneurons (Choy et al., 2017; Suzuki and Bekkers, 2011). To confirm that this synaptic organization was also true for Ntng1+ and Ntng1- cells, we injected Cre-dependent AAVs into Ntng1-Cre/Ai14 mice to express channelrhodopsin-2 (ChR2) in a focal subset of Ntng1+ cells (Figure S3a) and obtained whole-cell voltage-clamp recordings from uninfected Ntng1+ and Ntng1- cells away from the infection site (Figure S3c). We observed large, rapid inward currents in all recorded Ntng1- cells in response to brief (1 ms) light pulses but only small, if any, inward currents in Ntng1+ cells (Figures S3, d and e). Thus, the Ntng1-Cre mouse expresses Cre-recombinase in a subpopulation of PCx principal neurons whose laminar, morphological, intrinsic, and local connectivity properties are congruent with those of SLs. Therefore, henceforth we refer to Ntng1+ and Ntng1- cells within layer II of PCx as SLs and presumptive PYRs respectively.

Although SLs do not receive excitatory inputs from other SLs, we asked if they receive excitatory inputs from other PCx neurons. We focally injected nonconditional AAVs into Ntng1-Cre/Ai14 mice to express ChR2 in all PCx neurons (Figure S3f). We now observed even larger, rapid inward currents in PYRs compared to the responses in experiments using Cre-dependent AAVs (Figures S3, g and h), consistent with the dense recurrent connectivity between PYRs. Interestingly, we now observed small inward currents in SLs indicating that they do receive some intracortical excitatory inputs from other PCx neurons, although these were 5.4x weaker than inputs received by PYRs (Figures S3, f-g).

### SLs can be reliably identified in extracellular recordings of Ntng1-Cre mice

Having established that the Ntng1-Cre mouse line provides selective genetic access to SLs and confirmed key findings about the connectivity profiles of SLs and PYRs, we began probing functional differences between SLs and PYRs *in vivo*. We first injected Cre-dependent AAVs expressing the inhibitory opsin, Jaws (Chuong et al., 2014), into anterior PCx of Ntng1-Cre/Ai14 mice (Figure 1i). After allowing 3-4 weeks for Jaws expression, we recorded extracellular spiking activity of layer II PCx neurons at the injected site in awake, head-fixed mice. We used an optogenetic tagging strategy to classify recorded cells as either SLs or PYRs – we presented a series of brief 635 nm light pulses (1,000 pulses, 300 ms, 1 Hz), through either a 50 μm optic fiber attached to the recording probe or through a 200 μm optic fiber lowered separately to a distance of 0.5-0.8 mm from the recording probe, to suppress spiking in Ntng1+ cells (Figures 1, j and k). Cells were classified as SLs if their activity was reliably (i.e. spike rate greater in the baseline than in the light-on period, two-tailed t-test, p < 1×10^-7^) and rapidly (i.e. latency < 10 ms after light on, determined using change point analysis) suppressed by light (Rowland et al., 2018; Wolff et al., 2014). The remaining units were classified as PYRs. Across all experiments, approximately half of the recorded cells were SL based on these selection criteria (fraction of optogenetically identified SLs per experiment, mean ± s.d.: 0.486 ± 0.142; n = 15 experiments, 10 mice; Figure 1l). To minimize false negatives due to poor infection and/or mistargeted recordings, we omitted experiments in which less than 25% of cells were suppressed.

SLs had higher spontaneous spike rates than PYRs (median SL: 1.99 Hz, n = 426 cells; median PYR: 1.03 Hz, n = 464 cells; p = 1.02×10^-20^, unpaired t-test between means; Figure 1m), indicating that we can resolve differences in spiking activity between SLs and PYRs. Importantly, the overlapping distributions of SL and PYR spontaneous spike rates indicate that our opto-tagging protocol was not biased towards identifying cells with higher spike rates. Thus, the ability to record from PCx layer II principal neurons and reliably distinguish SL activity from PYR allowed us to next directly compare their odor responses.

### SLs are less odor selective and their population responses are less sparse than those in PYRs

We presented one-second-long pulses of odorized air and analyzed spiking activity aligned to the inhalation phase of the first full sniff after odor presentation (Bolding and Franks, 2017) (Figures 2, a and b). We first examined the distributions of activated and suppressed responses of SLs and PYRs, defined using a response index, where a value of 1 or −1 indicates that an observer can perfectly discriminate an increase or decrease in firing rate relative to baseline (pre-odor sniff) activity. Both cell types exhibited similar patterns of odor-activated and -suppressed responses, although SLs had slightly more suppressed odor responses than PYRs (Figures 2, c and d), likely due, in part, to their higher baseline firing rates.

**Figure 2.**
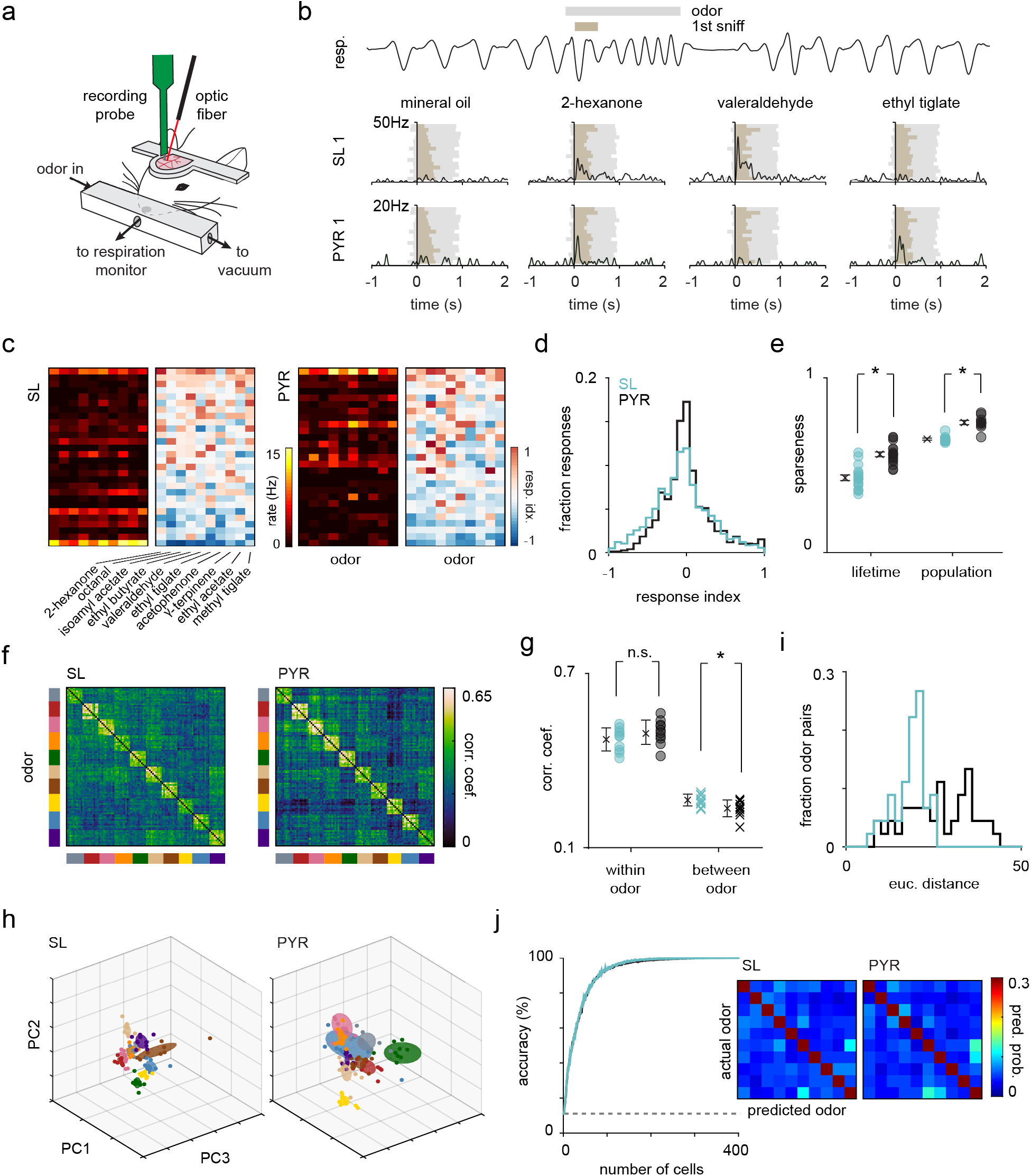
SLs and PYRs have distinct odor response properties. (**a**) Schematic of experimental setup for recording odor responses from awake, head-fixed Ntng1-Cre mice. (**b**) Example respiration trace from one odor trial (top). A 1-second long odor pulse (grey) is triggered on exhalation. Spiking activity is aligned to the onset of the subsequent inhalation (brown). Example SL and PYR responses across trials to different odors with spiking activity aligned to inhalation onset (bottom raster plots). Each tick indicates a spike. (**c**) Heatmaps of spike rates in the first sniff and response indices (area under the ROC curve x 2 - 1) for cell-odor pairs from an example experiment for SLs and PYRs shown separately. (**d**) Distribution of response indices for SL (blue, n=4260) and PYR (black, n=4640) cell-odor pairs across all experiments. (**e**) Lifetime sparseness for SLs (blue) and PYRs (black). Each circle is the average of all cells within an experiment (n=15 experiments). Population sparseness for SL and PYRs. Each circle is the average for an odor across all experiments (n=10 odors). (**f**) Trial-by-trial z-scored spike count correlation matrices sorted by odor for SLs and PYRs. Correlation matrices were generated for each experiment individually and then averaged across experiments. (**g**) Mean correlation across trials within each odor (circles) for SLs (blue) and PYRs (black) and between odors (crosses). (**h**) Principal components (PC) analysis was performed on z-scored SL (left) and PYR (right) pseudopopulation responses. Responses were projected onto the first three PCs. Each colored sphere represents an odor centered on the mean and encompasses one standard deviation. Colored dots indicate individual trials. (**i**) Distribution of pairwise Euclidean distances between the mean of the first 6 PCs for SLs (blue) and PYRs (black). (**j**) SL (blue) and PYR (black) odor classification accuracy determined using a linear support vector machine as a function of pseudopopulation size. The decoder was asked to classify responses (spike counts in a 500-ms time window beginning at odor inhalation) on a single trial to the panel of ten odors. Classification accuracy was averaged across 100 iterations for each pseudopopulation size. Confusion matrices, averaged across 100 iterations, on the right show the probability that an odor was predicted.

To more quantitatively analyze the response properties of SLs and PYRs, we determined their odor selectivity by calculating a lifetime sparseness value for each cell, which describes its firing rate distribution across the panel of odors. A value of 0 means the cell responded identically to all odors and a value of 1 means the cell responded to only one odor. SLs had significantly lower lifetime sparseness values than PYRs, indicating that SLs are less odor selective than PYRs (mean across experiments: SL, 0.424 ± 0.0162; PYR, 0.558 ± 0.0137; n = 15 experiments; p = 4.33×10^-7^, unpaired t-test; Figure 2e). We also calculated the population sparseness for SLs and PYRs, which describes the firing rate distribution of the population to each odor. Here, a value of 0 means all cells contributed equally to the population odor response and a value of 1 means the population odor response was driven by just one neuron. The SL population response to each odor was significantly less sparse than the PYR population response (mean across odors: SL, 0.686 ± 0.0027; PYR, 0.755 ± 0.0095; n = 10 odors; p = 1.61×10^-6^, unpaired t-test; Figure 2e). Together, these data indicate that SLs are less odor selective and less sparse than PYRs.

### Odors are slightly more discriminable in PYRs than in SLs

We then compared the reliability and discriminability of SL and PYR odor responses. We used single-trial population response vectors to generate trial-by-trial correlation matrices for SLs and PYRs (Figure 2f). We then calculated the mean correlation across trials within each odor as a measure of response reliability, and the mean correlation between different odors as a measure of odor discriminability. By this metric, response reliability was similar between SLs and PYRs (mean ± s.d.: SL, 0.474 ± 0.0396; PYR, 0.499 ± 0.0417; n = 10 odors; p = 0.191, unpaired t-test), but odors were slightly more discriminable in PYR responses (mean ± s.d.: SL, 0.266 ± 0.0201; PYR, 0.238 ± 0.0286; n = 45 odor pairs; p = 0.0193; Figure 2g).

To further investigate the discriminability of odor responses, we performed Principal Components Analysis (PCA) and, for visualization, projected SL and PYR responses onto their first three principal components for each odor (Figure 2h). We quantified the Euclidean distances between the means of the first six principal components of pairs of odors and found that again, odors were more separable in PYR responses (mean ± s.d.: SL, 18.3 ± 4.54; PYR, 26.8 ± 9.25; n = 45 odor pairs; p = 3.21×10^-7^, unpaired t-test; Figure 2i). Finally, we asked if a downstream decoder could more accurately identify odors based on PYR versus SL odor-evoked activity. We used a linear support vector machine trained and tested on single-trial spike count vectors to classify responses to the panel of ten odors. Odor classification accuracy, at all pseudopopulation sizes, was similar in SLs and PYRs (Figure 2j, left). To summarize, odors were more discriminable in PYRs than in SLs, consistent with the idea that the recurrent network decorrelates responses to different odors. However, the trial-by-trial variability and odor decoding performance of SLs and PYRs were equivalent.

Interestingly, we noticed some subtle but distinct off-diagonal structure in the correlation matrices for SLs and PYRs (Figure 2f). For example, valeraldehyde and γ-terpinene responses were more strongly decorrelated from other odor responses in PYRs than in SLs. Also, methyl tiglate responses were more similar to ethyl tiglate than acetophenone in PYRs, while the reverse was true in SLs. These differences were similarly reflected in the confusion matrices showing predicted odors when decoding using only 25 randomly selected cells for SLs and PYRs (scale adjusted to emphasize classification errors) (Figure 2j, right). If both SL and PYR responses merely reflected their sensory inputs, then similar odors would evoke similar response patterns in both cell types and misclassifications would be independent of cell type. Instead, our data suggest that SLs and PYRs are extracting and processing odor information differently.

### SLs and PYRs have similar response latencies

Next, we determined if odor-evoked activity in SLs precedes that in PYRs. We measured the time from inhalation onset to both the response peak (Figure 3a) and response onset (i.e. when the odor-evoked spike rate exceeded two standard deviations above baseline activity) (Figure 3c) in a 100 ms time window after inhalation onset for each cell-odor pair. Only activated responses (response index > 0) were analyzed. In both measures, the distributions of response latencies of SLs and PYRs were similar (median peak latency: SL, 59.7 ms; PYR, 58.7 ms; Wilcoxon rank-sum test, p = 0.0293; median onset latency: SL, 35.0 ms; PYR, 36.2 ms; Wilcoxon rank-sum test, p = 0.831, Figures 3, b and d). Furthermore, the performance of SLs and PYRs in a classifier trained on spike counts in a 10- or 30-ms sliding window beginning at inhalation onset were equivalent (Figure 3e). This indicates that odor information accumulates at the same rate in SLs and PYRs. Taken together, our data do not appear to support a sequential activation model in which SL activity precedes PYR activity. However, we cannot rule this out entirely based on our analyses of response latencies (see Discussion).

**Figure 3.**
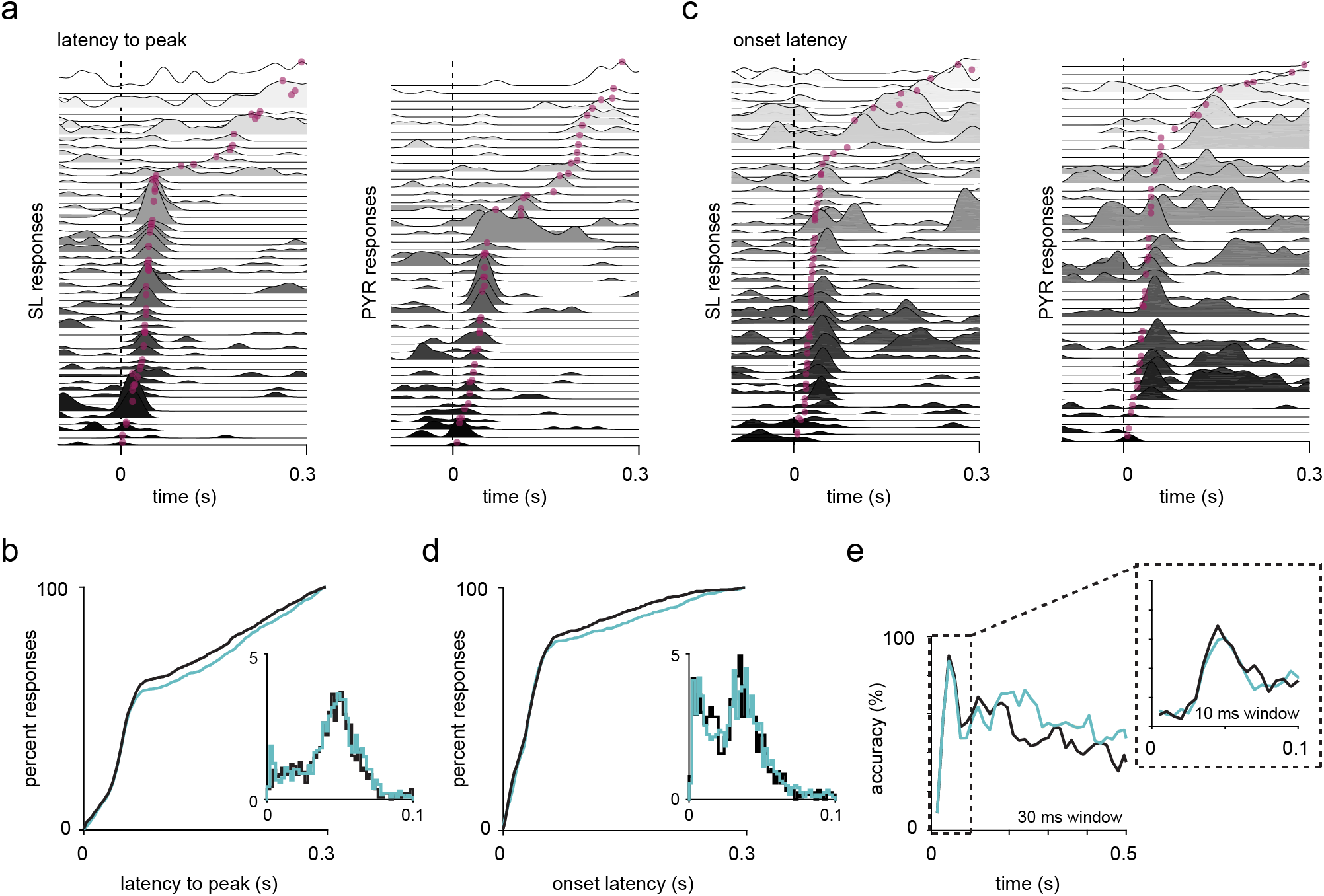
SLs and PYRs have similar response latencies. (**a**) SL and PYR odor responses from one example experiment sorted by latency to response peak. Dotted line at t=0s marks inhalation onset. Magenta dots indicate the time of response peaks. (**b**) Cumulative distribution of latencies to peak for active (response index > 1) SL (blue, n=1408) and PYR (black, n=1368) cell-odor pairs. Inset shows histogram of latencies to peak within 100ms of inhalation onset. (**c**) As in (**a**) but sorted by onset latencies. Magenta dots indicate the time at which each odor response reaches 2 s.d. above pre-odor baseline. (**d**) As in (**b**) but showing onset latencies. SL (blue, n=1275) and PYR (black, n=1265) cell-odor pairs. (**e**) Odor classification accuracy using a linear support vector machine in a sliding window of either 30ms or 10ms (inset).

### Optogenetic suppression of SLs does not weaken or reshape PYR odor responses

We next asked if SL activity is required to drive odor responses in PYRs. We optogenetically suppressed SLs while recording baseline and odor-evoked activity in PYRs, thus isolating the effect of removing SL input on PYRs (Figure 4a). To maximize the area of PCx that we suppressed, we injected Cre-dependent Jaws in three sites along the anterior-posterior axis of PCx (Figure 4b) and we used a thicker (400 μm) optic fiber, positioned at an angle to the recording probe (as in Figure 1b). We opto-tagged SLs at the end of each experiment (fraction of identified SLs, mean ± s.d.: 0.432 ± 0.111, n = 8 experiments, 6 mice). We presented a panel of five monomolecular odorants, and on alternating odor trials, delivered a three-second-long light pulse spanning approximately one second before to one second after odor presentation. Odor delivery was triggered at the onset of exhalation, introducing a little jitter (~100 ms) in the amount of time the light was on before the odor was delivered. During a 500 ms-period before odor onset, we observed strong and sustained light-evoked suppression of SLs, as well as a slight decrease in the firing rate of PYRs, indicating that SL activity, at baseline, contributes to spontaneous PYR activity (Figure 4c).

**Figure 4.**
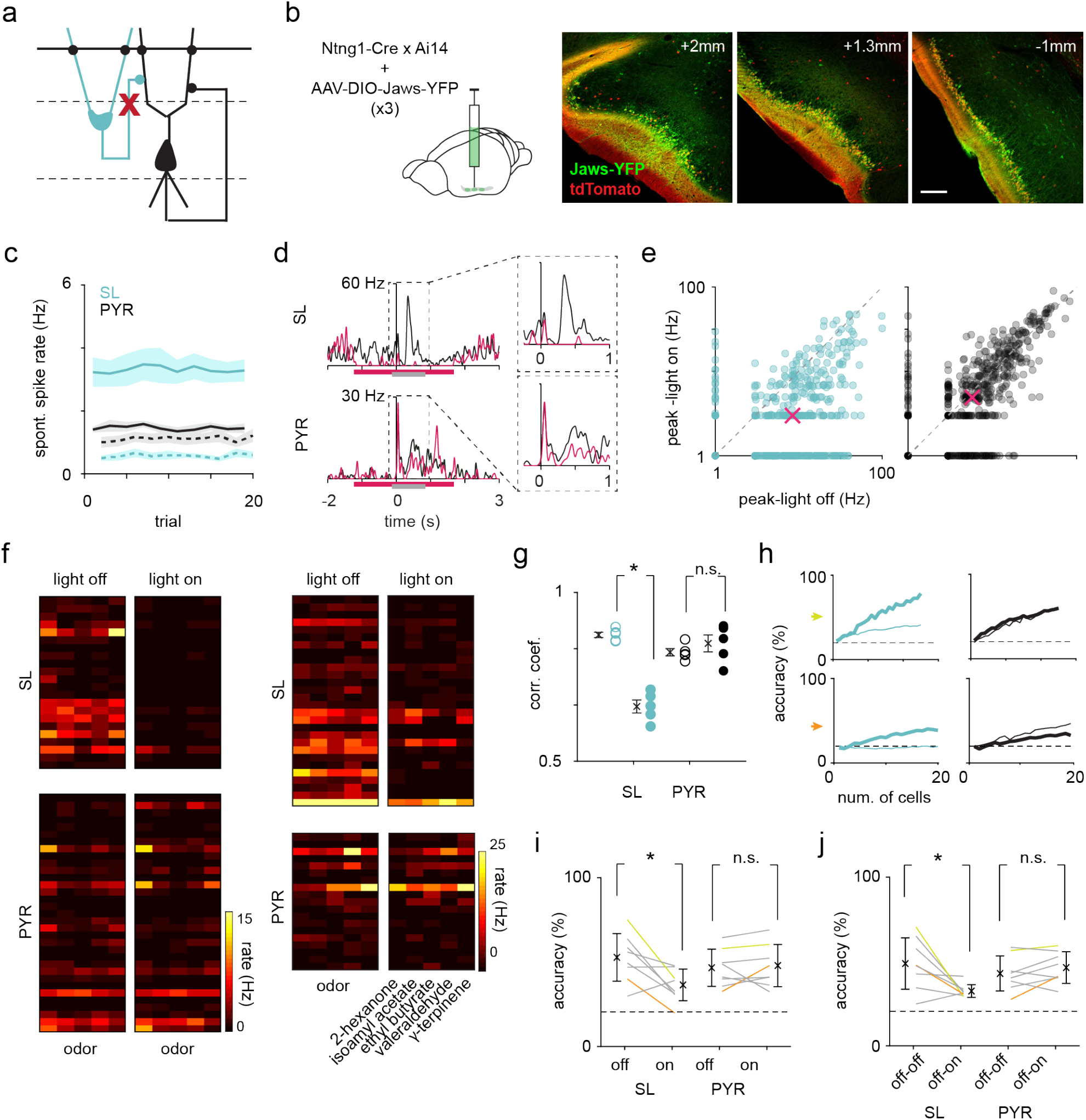
Optogenetic suppression of SLs does not weaken or reshape PYR odor responses. (**a**) Experimental setup. SLs were optogenetically suppressed during recording. (**b**) Histology showing Jaws expressed selectively in SLs. Jaws was injected in 3 sites along the anterior-posterior axis (left to right) of PCx to maximize SL suppression. Scale bar is 200 μm. (**c**) Average spontaneous spike rate of SLs (blue) and PYRs (black) during *light-off* (solid line) and *light-on* (dotted line) trials. (**d**) Example SL and PYR odor responses during alternating *light-on* (pink trace) and *light-off* (black trace) trials. Responses are aligned to inhalation onset and are shown as trial-averaged PSTHs. The pink bar indicates when the light was on and the grey bar indicates when the odor was presented. (**e**) Odor-evoked response peaks for activated SLs (left, n=439 cell-odor pairs) and PYRs (right, n=603 cellodor pairs) during *light-on* trials plotted against *light-off trials*. Median response peak is shown in pink. (**f**) Heatmaps of spike rates in the first sniff for SL and PYR cell-odor pairs from two example experiments, one with complete (left) and one with incomplete SL suppression (right). *Light-on* and *light-off* trials are plotted separately. (**g**) Correlation coefficients for response vectors in the light-on and light-off conditions (filled circles, n=5 odors) and in just the light-off condition (open circles, n=5 odors) for each odor. (**h**) Odor classification accuracy for SLs (blue) and PYRs (black) for two example experiments as a function of cell number in the *light-on* (thick line) and *light-off* (thin line) conditions separately (decoding within condition). (**i**) Summary plot for within condition decoding (yellow and orange lines represent example experiments in h). Each line is an experiment. (**j**) Odor classification accuracy for SLs and PYRs within each experiment, where the classifier was trained on trials from the *light-off* condition and tested on trials from the *light-on* condition (decoding across condition). Each line is an experiment.

We then examined the odor-evoked spike rates of SLs and PYRs. To avoid confounds of odor-evoked suppression, we only considered activated responses (response index > 0 during *light-off* trials). During the *light-on* trials, odor-evoked activity, measured as peak firing rate in the first sniff, was markedly suppressed in SLs with no significant effect on PYRs (median response peak: SL, *light-off*: 7.43 Hz, *light-on*: 1.99 Hz; n = 439 cell-odor pairs; paired t-test between means, p = 2.61×10^-31^; PYR, *light-off*: 4.79 Hz, *light-on*: 3.90 Hz; n = 603 cell-odor pairs; paired t-test between means, p = 0.208; Figures 4, d-f).

Although the strength of individual PYR responses was only minimally affected by SL suppression, the population response patterns may have changed. To quantify changes in odor representations between conditions, we calculated the correlation between the trial-averaged population response for each odor in the *light-on* and the *light-off* conditions. As a control, we calculated the correlation between the trial-averaged population response for each odor in alternating *light-off* trials. SL response patterns across odors were significantly altered in the *light-on* trials compared to the *light-off* trials (*off-off*, 0.870 ± 0.0079; *on-off*, 0.657 ± 0.0193; unpaired t-test, p = 7.40×10^-6^; Figure 4g), while PYR response patterns were unchanged (*off-off*, 0.818 ± 0.0107; *on-off*, 0.844 ± 0.0250; unpaired t-test, p = 0.365; Figure 4g).

Furthermore, odor identity decoding analyses revealed a substantial decrease in SL performance in the *light-on* condition across all experiments, with no change in PYR performance (Figures 4, h and i). In fact, in one experiment where SL suppression was complete and decoding performance decreased to chance in the *light-on* condition, decoding in PYRs improved, perhaps due to an increase in the signal-to-noise ratio in the absence of SL input (Figure 4h, bottom). This improvement in PYR performance argues against the possibility that we did not see a substantial impairment in PYR responses because we were not suppressing a large enough number of SLs. These data indicate that PYRs can accurately represent odor identity when SLs are suppressed, but they do not tell us whether PYR representations remain unchanged. To address this question, we performed a decoding analysis across conditions, where the classifier was trained on *light-off* trials and then tested on *light-on* trials. This analysis also showed a substantial decrease in SL performance but, again, no change in PYR performance (Figure 4j), indicating that odor-evoked population activity in PYRs is independent of SL input.

### Chemogenetic suppression of SLs does not weaken or reshape PYR odor responses

Mouse PCx is a large structure measuring ~4 mm along its anterior-posterior axis with long-range intracortical connectivity (Franks et al., 2011; Hagiwara et al., 2012; Johnson et al., 2000). To maximize the number of SLs that we suppressed, we repeated the SL suppression experiment using a chemogenetic approach (Figure 5a), where AAVs expressing a Cre-dependent inhibitory DREADD, hm4di was injected at four sites along the anterior-posterior axis of PCx. At one of the sites, we co-injected AAVs expressing Cre-dependent Archaerhodopsin (Arch) to opto-tag SLs (fraction of identified SLs, mean ± s.d.: 0.546 ± 0.146; n = 5 experiments, 4 mice; Figure 5b). The DREADD agonists, CNO or Compound 21 (C21), were administered intraperitoneally during the recording. Following CNO/C21 administration, we observed a gradual decrease in SL spontaneous activity and a small decrease in PYR spontaneous activity (Figure 5c).

**Figure 5.**
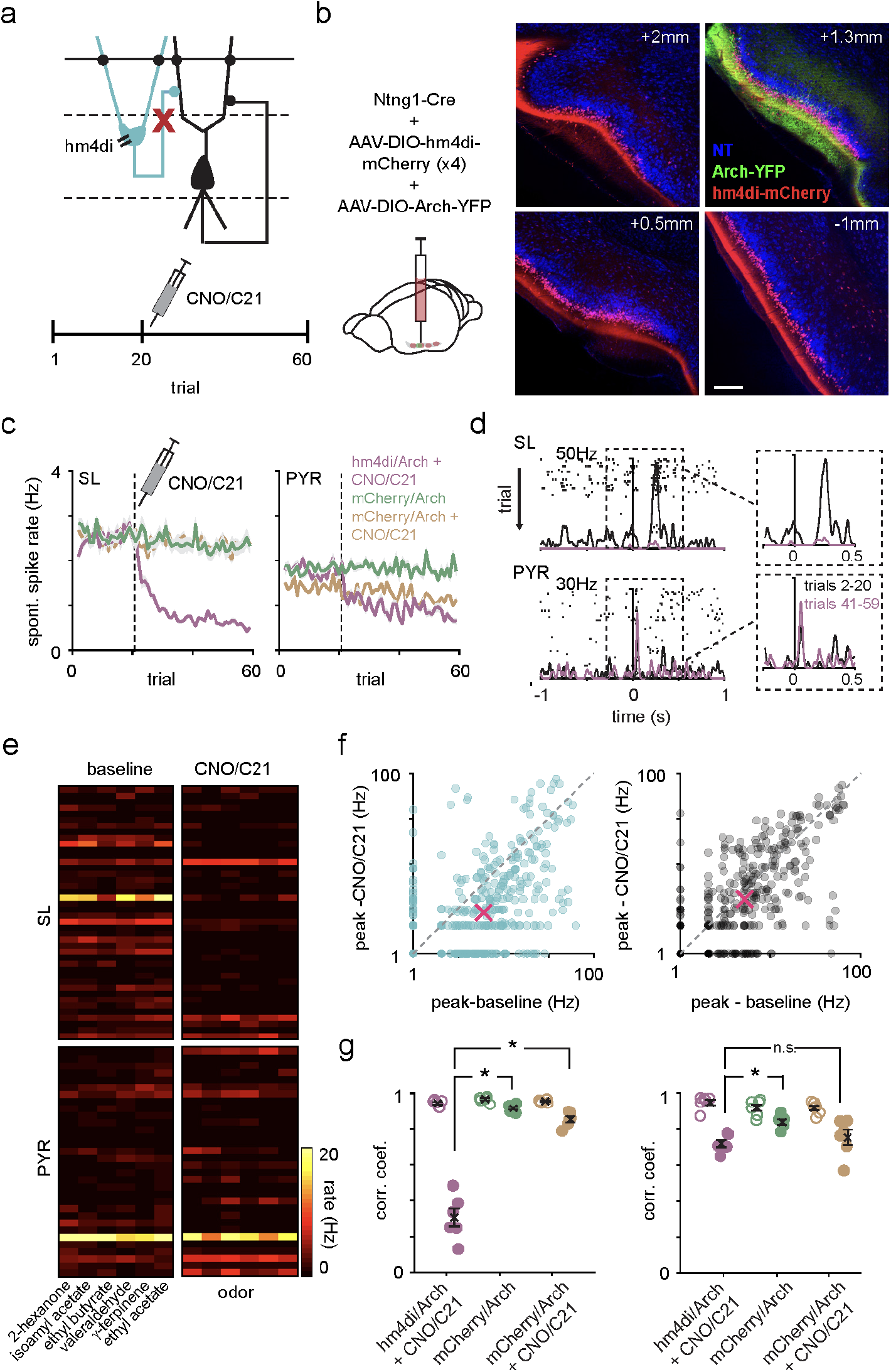
Chemogenetic suppression of SLs does not weaken or reshape PYR odor responses. (**a**) Top: schematic showing the point of perturbation of the circuit in PCx. SLs are selectively suppressed by activation of hm4di during recording. Bottom: CNO/C21 is administered intraperitoneally after trial 20. (**b**) Coronal sections showing selective expression of hm4di-mCherry in SLs along the length of PCx and Arch. AAVs carrying hm4di or mCherry (control experiments) was injected in 4 sites along the anterior-posterior axis (left to right) of PCx to maximize SL suppression. Arch was co-injected in one of the sites for opto-tagging. Scale bar is 200 μm. (**c**) Average spontaneous spike rate of all SLs (left) and PYRs (right) as a function of trials from experimental and control groups. CNO/C21 is administered intraperitoneally after trial 20 (dotted line). (**d**) Odor response of an example SL and PYR from a hm4di/Arch+CNO/C21 mouse across trials. CNO/C21 trials are shown in pink and baseline trials in black. (**e**) Heatmaps of spike rates in the first sniff for SL and PYR cell-odor pairs from two example experiments with baseline and CNO/C21 trials plotted separately. (**f**) Odor-evoked response peaks for activated SLs (left, n=391 cell-odor pairs) and PYRs (right, n=348 cell-odor pairs) during CNO/C21 trials plotted against baseline trials. Median response peak is shown in pink. (**g**) Correlation coefficients of pseudopopulation response vectors of baseline and CNO/C21 trials (filled circles, n=6 odors) or alternating baseline trials (open circles, n=6 odors) for each odor for SLs (left) and PYRs (right) for experimental and control groups.

We examined SL and PYR odor-evoked spike rates before (trials 2-20) and after (trials 41-59) hm4di-mediated suppression of SLs, again only considering cell-odor pairs with activated responses before CNO/C21 administration. As in the optogenetic suppression experiments, the odor-evoked response peaks in SLs were substantially lower after CNO/C21 injection, while the odor-evoked response peaks in PYRs were only slightly diminished (median response peak.: SL, baseline: 5.05 Hz, CNO/C21: 1.84 Hz; n = 391 cell-odor pairs; paired t-test between means, p = 2.44×10^-5^; PYR, baseline: 4.18 Hz, CNO/C21: 3.02 Hz; n = 348 cell-odor pairs; paired t-test between means, p = 0.0506; Figures 5, d-f).

To determine if the population odor responses in PYRs were altered after hm4di-mediated suppression of SLs, we calculated the correlation between the trial-averaged population response of SLs and PYRs for each odor before and after CNO/C21 injection. As a control, we calculated the correlation between the trial-averaged population response for each odor in alternating baseline trials. Responses in SLs changed markedly before and after CNO/C21 (baseline-baseline, 0.938 ± 0.0078; CNO/C21-baseline, 0.300 ± 0.0492; p = 1.59×10^-7^; Figure 5g). PYR responses also changed after CNO/C21 administration (baseline-baseline, 0.945 ± 0.0159; CNO/C21-baseline, 0.729 ± 0.0173; p = 2.20×10^-6^; Figure 5g), although this difference was much smaller than that observed in SLs (CNO/C21-baseline for SLs versus PYRs, p = 9.34×10^-6^). However, unlike the optogenetic suppression experiments, in which we compared alternating *light-on* and *light-off* trials, the chemogenetic suppression experiments required comparing responses before and after CNO/C21 had taken effect, which was typically 30 minutes and 20 trials later.

Therefore, to determine the extent to which the changes we observed after CNO/C21 administration in both SLs and PYRs were directly due to the suppression of SL activity by the activation of hm4di, rather than adaptation, desensitization, attentional changes, or repeated odor exposure (Jacobson et al., 2018), we recorded from Ntng1-Cre mice that were injected with Cre-dependent mCherry and Arch, for 60 trials without injecting CNO or C21 (fraction of identified SLs, mean ± s.d.: 0.449 ± 0.146, n = 5 experiments, 4 mice). Interestingly, we found that both SL and PYR responses in the early (trials 2-20) and later trials (trials 41-59) were not completely correlated, indicating that some changes in response patterns may occur with time or following repeated odor presentations (SLs, baseline-baseline, 0.953 ± 0.0052; CNO/C21-baseline, 0.913 ± 0.0089; p = 0.0028; PYRs, baseline-baseline, 0.923 ± 0.0063; CNO/C21-baseline, 0.841 ± 0.0149; p = 3.01×10^-4^; Figure 5g). Additionally, in a separate set of mice injected with AAVs expressing mCherry and Arch, we administered CNO/C21 to account for any non-specific, hm4di-independent effects of CNO/C21 (fraction of identified SLs, mean ± s.d.: 0.545 ± 0.229, n = 3 experiments, 3 mice). Similar to the previous control, we found that response patterns had changed between the baseline and CNO/C21 conditions in both SLs and PYRs (SLs, baseline-baseline, 0.945 ± 0.0059; CNO/C21-baseline, 0.852 ± 0.0162; p = 3.01×10^-4^; PYRs, baseline-baseline, 0.922 ± 0.0137; CNO/C21-baseline, 0.747 ± 0.0479; p = 0.0056; Figure 5g).

In summary, SL responses were substantially degraded following hm4di activation while PYR responses were only minimally affected. Importantly, the changes in PYR responses in the experimental group were comparable to the changes in PYR responses in the two control groups, indicating that the effect we observed in PYRs was not due to hm4di-mediated suppression of SLs (CNO/C21-baseline for PYRs in hm4di/Arch+CNO vs. mCherry/Arch+CNO, p = 0.731; Figure 5g). Thus, consistent with the optogenetic suppression experiments, PYR odor responses are largely independent of SLs.

## DISCUSSION

The Ntng1-Cre mouse line allowed us to distinguish between SLs and PYRs and selectively suppress SLs in extracellular recordings of layer II PCx neurons *in vivo*.We found that SLs were more broadly tuned, less sparse, and their population odor responses less discriminable compared to PYRs. However, these differences were relatively small. We did not detect a difference in odor response latencies between SLs and PYRs, and suppressing SLs, both optogenetically and chemogenetically, during odor presentation did not weaken or alter the response pattern of PYRs. Our data therefore do not support a two-step sequential model of activation in PCx, but rather suggest that these two cell types respond similarly and operate in parallel.

### Stimulus selectivity, discriminability, and response variability

Individual PCx neurons integrate convergent inputs from different combinations of glomeruli (Apicella et al., 2010; Davison and Ehlers, 2011). SLs almost exclusively receive afferent OB inputs, while PYRs receive both afferent OB inputs and dense intracortical inputs from SLs and other PYRs (Franks et al., 2011; Hagiwara et al., 2012; Johnson et al., 2000; Suzuki and Bekkers, 2011). If intracortical connectivity is random, then information about active glomerular combinations received by PCx neurons would be uniformly redistributed among PYRs via the recurrent network. Individual PYRs would therefore effectively sample a far larger subset of glomeruli than individual SLs, resulting in more broadly tuned and overlapping responses among PYRs (Suzuki and Bekkers, 2011). Alternatively, coordinated activity within PCx could actively sculpt the recurrent circuitry so that PYRs that are regularly co-activated, because they respond to similar chemical features, become preferentially interconnected resulting in more narrowly tuned, reliable, and discriminable responses than those in SLs. Our results are largely consistent with the latter model in that we found PYR responses to be more narrowly tuned and discriminable. A recent study showed that odor responses in layer III of PCx, which only contains PYRs, were more correlated between chemically similar odor stimuli than responses in layer II, which contain both SLs and PYRs, or the OB (Pashkovski et al., 2020), providing further evidence that associative inputs are structured and nonrandom. It was therefore surprising that PYR responses were not more reliable than SL responses.

Our findings on the stimulus selectivity and discriminability of SLs and PYRs are comparable to those reported in two zebrafish odor centers downstream of the OB: cells in the ventral telencephalon (Vv) receive primarily OB inputs and exhibit broader stimulus tuning and more overlapping odor representations than those in the dorsal telencephalon (Dp), which is the teleost homolog of PCx (Yaksi et al., 2009). Finally, in the control experiments for the DREADD suppression of SLs, we noticed that PYR responses changed more with time and/or stimulus presentation than SL responses. This result is consistent with an activity-dependent sculpting of recurrent connectivity so that PYR responses evolve while SL responses continue to faithfully represent the stimulus (Bolding et al., 2020; Jacobson et al., 2018; Pashkovski et al., 2020; Schoonover et al., 2021).

### Interpreting effect size

Although the response properties of SLs and PYRs are consistent with a more sensory and a more associative network respectively, the differences we observed were less pronounced than we expected. There are several possible explanations for this. First, although SLs and PYRs are the main two classes of excitatory principal neurons in Layer II, some neurons have intermediate morphological and intrinsic properties that lie on a gradient from canonical SLs to canonical PYRs (Suzuki and Bekkers, 2011; Wiegand et al., 2011; Yang et al., 2004). These cells, which may or may not be Ntng1+, could be diluting the differences that we observed. Second, although SLs receive stronger OB inputs than PYRs, this difference is relative and does not mean that OB inputs onto PYRs are weak or that PYRs are not directly driven by the OB, as this is sometimes interpreted. Third, unlike in slice recordings, the recurrent network and top-down inputs are preserved in our *in vivo* recordings. Therefore, while SL activity is driven largely by sensory input, PYR activity is also influenced by ongoing background activity that provide non-olfactory inputs to PYRs, rendering their odor representations more variable and less discriminable than perhaps expected (Stringer et al., 2019). Why did we find no difference in the response latencies of SLs and PYRs even though SLs responded approximately 2-4 ms earlier than PYRs in response to LOT stimulation in brain slice experiments or in anesthetized rats (Ketchum and Haberly, 1993; Suzuki and Bekkers, 2011; Wiegand et al., 2011)? *In vivo*, ongoing recurrent and top-down inputs in awake animals may depolarize PYRs and thereby shorten their response latencies. More importantly, electrical LOT stimulation generates a single, synchronous barrage of afferent inputs into PCx. However, different mitral cells respond to odors with variable latencies that tile the entire ~300 ms sniff cycle (Bolding and Franks, 2018; Cury and Uchida, 2010; Shusterman et al., 2011), making it almost impossible to compare SL and PYR onset latencies to a given odor-activated input. Precise optogenetic stimulation of OB glomeruli with high temporal precision (Chong et al., 2020) can be used to address this question more directly.

### Potential limitations of optogenetic and chemogenetic suppression

With optogenetics, we were able to effectively suppress the SLs we were recording from on alternating trials, thus providing a robust comparison of PYR responses with and without SL input. However, PCx spans 3-4 mm along its anterior-posterior axis and is a long-range recurrent network with neurons forming intracortical connections with neurons >1 mm away (Franks et al., 2011; Hagiwara et al., 2012; Johnson et al., 2000). Hence, to effectively remove SL inputs onto PYRs, we needed to suppress SLs located >1 mm from the recording site. However, the spatial resolution of optogenetic suppression using 532 nm light, determined using a photobleaching assay, is slightly less than 1 mm (Li et al., 2019). To broaden the area of suppression, we used a thicker (400μm) optic fiber and Jaws, a red-light (635 nm) activated opsin, as red light scatters less in tissue (Chuong et al., 2014; Wiegert et al., 2017). We injected the virus in 3-4 sites along the anterior-posterior axis of PCx to suppress axon terminals, at the recording site, of cell bodies that were outside of the light cone. Nevertheless, we do not know how effectively we suppressed SLs far away from the recording site, which was also the center of illumination. Chemogenetics alleviates the spatial limitations of optogenetics, but the suppression of odor-evoked activity among SLs was less robust than in the optogenetic experiments. Additionally, we had to wait 20-30 minutes for cells to be suppressed which allowed for shifts in odor representations. Novel techniques that enable more effective axon terminal suppression with high spatiotemporal precision and efficacy (Mahn et al., 2021) may enable a more robust interrogation of the effects of suppressing SLs on PYRs. These caveats notwithstanding, both the optogenetic and chemogenetic experiments showed effective SL suppression and indicated that SL suppression had little to no effect on PYR odor responses.

### Roles of SLs and PYRs in odor processing

Our data do not support a two-stage sequential activation model, where odor information from the OB is transmitted first to SLs and then from SLs to PYRs. Instead, both SLs and PYRs can be driven by OB input and appear to differentially process odor information, as suggested by the differences we observed in the off-diagonal structure of their correlation matrices. The distinct afferent versus associative connectivity between SLs and PYRs perhaps enables each cell type to extract different features of odor information and route this information to distinct downstream regions (Diodato et al., 2016; Mazo et al., 2017). For example, SLs preferentially project to the posterior PCx, posteromedial cortical amygdala and lateral entorhinal cortex while PYRs project back to the OB, medial prefrontal and orbitofrontal cortices.

Although it may not be surprising that PYRs are able to respond to odors without SL input, our finding that PYR odor representations are essentially equivalent with and without SL input is nevertheless puzzling. Why might this be, and what does this imply about the function of SL projections onto PYRs? Inputs onto PYRs from other PYRs are approximately five times stronger than inputs from SLs (Figure 2e). It might be that under our very simplified conditions, with pure odors delivered at relatively high concentrations (0.3% vol./vol.), the contribution from SLs becomes insignificant and may be more apparent in more naturalistic odor environments. Alternatively, in the optogenetic experiments, we suppressed SLs hundreds of milliseconds before odor onset, which could be sufficient time for the network to re-equilibrate (Bolding and Franks, 2018). Perhaps rapid suppression of SLs during or immediately before the odor would have a more pronounced effect. We also presented odors passively, in non-behaving mice, and it is possible that differences between, and dependencies on, the different cell types might emerge in mice engaged in a behavioral task (Chen et al., 2015). However, this seems unlikely given recent evidence that odor representations in PCx do not appear to encode odor value (Gadziola et al., 2020; Millman and Murthy, 2020; Wang et al., 2020) and that any ‘top-down’ effects would almost certainly influence PYR output more than SL. Finally, perhaps SLs and PYRs do not simply represent odor identity and we are looking at the wrong feature of the odor response. Future experiments will be required to resolve this issue, but our data make clear that odor processing in PCx does not occur through a two-stage process mediated first by SLs and then by PYRs.

### Comparing PCx to the hippocampus

PCx and the hippocampus are analogous structures. Both are trilaminar paleocortices with extensive recurrent networks (Guzman et al., 2016) that support unsupervised learning (Barkai et al., 1994; Haberly, 2001; Rolls, 2013). In terms of morphology and connectivity, PYRs are analogous to CA3 pyramidal cells and SLs to DG granule cells. The roles of DG and CA3 in the encoding of episodic memories are relatively well characterized; DG granule cells are thought to drive the acquisition of memories, but not retrieval, and perform pattern separation while CA3 pyramidal cells are thought to store and retrieve memories and perform pattern completion (Hainmueller and Bartos, 2020; Leutgeb et al., 2007; Madar et al., 2019; Neunuebel and Knierim, 2014). It is important to note that although DG granule cells synapse onto CA3 pyramidal neurons, the DG-CA3 circuit is not exclusively sequential. CA3 also receives direct input from the entorhinal cortex, and when DG projections to CA3 are silenced or ablated, the ability of CA3 to retrieve memories is unimpaired (Lee and Kesner, 2004). In our previous study with the Ntng1-Cre mouse line, we found that PYRs were more robust to degraded afferent OB inputs and exhibited more persistent odor-evoked activity after stimulus offset than SLs (Bolding et al., 2020). Both of these features are consistent with an associative network performing computations that stabilize and enable reliable retrieval of odor ensembles, characteristic of a pattern completing network (Inagaki et al., 2019; Rolls, 2013). However, there is yet to be evidence implicating SLs, specifically, as pattern separators in the olfactory circuit (Chapuis and Wilson, 2011). A key property of DG granule cells that enable them to effectively decorrelate similar activity patterns is their sparseness (Leutgeb et al., 2007). However, we found that SLs had higher spontaneous spike rates, were individually more broadly tuned, and their population response less sparse than PYRs.

### Conclusion

In all, our data do not support a sequential processing model for PCx in which odor information from OB is first integrated in SLs, and then routed from SLs to PYRs. Instead, we propose that these distinct excitatory cell types in PCx form largely parallel processing schemes, where each cell type receives varying amounts of sensory and associative inputs, performs distinct local computations and projects the information to distinct downstream regions. SLs and PYRs both receive afferent OB inputs. However, because SLs are less driven by intracortical inputs, downstream regions receiving SL inputs may receive odor information that more closely reflects the chemistry of the stimulus and are more sensitive to changes in afferent inputs. On the other hand, PYRs may serve as a storage for stable odor representations, that are robust to variable stimulus conditions.

## MATERIALS AND METHODS

All experimental protocols were approved by Duke University Institutional Animal Care and Use Committee. Information about the Ntng1-Cre mouse line is described in Bolding et al., 2020. Methods for in vivo extracellular recording and preliminary analysis have been reported in detail in Bolding and Franks, 2017 and Bolding et al., 2020, and are summarized here. All data are shown as mean ± sem, unless otherwise stated.

### Subjects

All experiments were performed on adult Ntng1-Cre knock-in mice, which were generated by inserting the DNA sequence encoding Cre-recombinase at the start codon of the Ntng1 gene using CRISPR. All mice were heterozygotes, crossed to either Ai14 (B6.Cg-Gt(ROSA)26Sor^tm14(CAG-tdTomato)Hze^/J, 007914) or C57BL/6J (000664) mice obtained from The Jackson Laboratory.

### Stereotaxic injections and headpost placement

Mice were given Buprenorphine-SR (0.1mg/kg, s.c.) ten minutes before the start of surgery and then placed in a closed glass box filled with 5% isoflurane to induce anesthesia. Afterwards, they were moved to a stereotaxic frame and maintained on isoflurane anesthesia (0.5-2% in 0.8L/min O_2_) throughout the procedure. A local anesthetic, Bupivacaine, was injected under the skin over the skull and an incision was made along the midline to expose the skull. Four burr holes were made over piriform cortex using coordinates relative to bregma and adeno-associated viruses (either AAV8-hSyn-DIO-Jaws-eYFP or AAV5-hSyn-DIO-ArchT3.0-eYFP or AAV5-hSyn-DIO-hm4di-mCherry or AAV5-hSyn-FLEX-mCherry, all obtained from UNC Vector Core) were then delivered through a glass pipette using a Nanoject pump (Drummond) (200nL/site at 90 nl/min). The following coordinates were used (in mm, AP 2.0, ML 2.15, DV 3.8; AP 1.3, ML 2.82, DV 3.95; AP 0.5, ML 3.5, DV 4.05; AP −0.6, ML 3.8, DV 4.10). After slowly retracting the pipette, burr holes were covered using KwikCast sealant and a custom titanium headpost was lowered onto the exposed skull and attached using Metabond (Parkell, Inc.). Mice were placed on a heating pad for 30 minutes after surgery while they recovered from anesthesia.

### *In vitro* electrophysiology and analysis

Mice were anesthetized with isoflurane and decapitated, and the cortex was quickly removed in ice-cold artificial CSF (aCSF). Parasagittal brain slices (300 μm) were cut using a vibrating microtome (Leica) in a solution containing (in mM): 10 NaCl, 2.5 KCl, 0.5 CaCl2, 7 MgSO4, 1.25 NaH2PO4, 25 NaHCO3, 10 glucose, and 195 sucrose, equilibrated with 95% O2 and 5% CO2. Slices were incubated at 34°C for 30 min in aCSF containing: 125 mM NaCl, 2.5 mM KCl, 1.25 mM NaH2PO4, 25 mM NaHCO3, 25 mM glucose, 2 mM CaCl2, 1 mM MgCl2, 2 NaPyruvate. Slices were then maintained at room temperature until they were transferred to a recording chamber on an upright microscope (Olympus) equipped with a 40x objective.

For current clamp recordings, patch electrodes (3-6 megohm) contained: 130 Kmethylsulfonate, 5 mM NaCl, 10 HEPES, 12 phosphocreatine, 3 MgATP, 0.2 NaGTP, 0.1 EGTA, 0.05 AlexaFluor 594 cadaverine. For voltage-clamp experiments, electrodes contained: 130 D-Gluconic acid, 130 CsOH, 5 mM NaCl, 10 HEPES, 12 phosphocreatine, 3 MgATP, 0.2 NaGTP, 10 EGTA, 0.05 AlexaFluor 594 cadaverine. Voltage- and current-clamp responses were recorded with a Multiclamp 700B amplifier, filtered at 2-4 kHz, and digitized at 10 kHz (Digidata 1440). Series resistance was typically ~10 megohm, always <20 megohm, and was compensated at 80%–95%. The bridge was balanced using the automated Multiclamp function in current clamp recordings. Data were collected and analyzed off-line using AxographX and IGOR Pro (Wavemetrics). Junction potentials were not corrected. Voltage-clamp responses were recorded with a Multiclamp 700B amplifier, and digitized at 10 kHz (Digidata 1440); evoked responses were low-pass filtered at 4 kHz.

#### Intrinsic properties

Recordings were targeted to SLs (tdTomato+) and PYRs (unlabeled) located in layer II. Cells were also visualized using a fluorescent indicator (Alexa 594 Cadaverine) to confirm that they were correctly identified as SL or PYR. Intrinsic properties were measured at current clamp in response to a series of 1-second-long current pulses stepped up in 50 pA increments.

#### Synaptic connectivity

All experiments were performed 3-4 weeks after virus injection (AAV-hSyn-DIO-Chr2 or AAV-hSyn-ChR2). We recorded from uninfected SLs (tdTomato+) and PYRs (unlabeled) adjacent to the infection site. To ensure cells were uninfected we first examined responses to weak, 1-second long light pulses (470 nm, CoolLED) delivered through the 40x objective. Cells that exhibited large and sustained photocurrents were discarded. Uninfected cells were held at either −70 mV or +5 mV to isolate excitatory or inhibitory synaptic currents, respectively. Brief (1 ms, ~10 mW) pulses were delivered every 10 seconds to activate ChR2+ axon terminals.

### *In vivo* extracellular recordings

#### Electrode placement and data acquisition

Recordings were performed 3-6 weeks after virus injections. On the day of the recording, a craniotomy was made around the burr hole to enlarge the opening and dura mater was removed. A 32-site polytrode acute probe, either with a 50μm optic fiber attached (A1×32-Poly3-5mm-25s-177-OA32LP, Neuronexus, Ann Arbor, MI) or without (A1×32-Poly3-5mm-25s-177-A32, Neuronexus) was positioned over the craniotomy and lowered into one of the virus injection sites using a Patchstar Micromanipulator (Scientifica, UK). Recordings were targeted to 3.8mm-4.3mm ventral to the brain surface and the probe was lowered until a band of intense spiking activity, reflecting the densely packed layer II of piriform cortex, was observed. Electrophysiological signals were acquired through an A32-OM32 adaptor (Neuronexus), digitized at 30kHz at a Cereplex digital headstage (Blackrock Microsystems, Salt Lake City, UT) and recorded using a Cerebus multichannel data acquisition system (Blackrock Microsystems). Experimental events (e.g. odor delivery times, laser pulse times) and respiration signals were acquired at 2kHz by analog inputs of the Cerebus system. Respiration was monitored using a microbridge mass airflow sensor (Honeywell AWM3300V) positioned opposite the animal’s nose.

#### Spike sorting

Individual units were isolated using Spyking-Circus (https://github.com/spyking-circus, Yger et al., 2018). Isolated clusters were manually curated to remove those with >1% of ISIs violating the refractory period (<2 ms) or that looked like noise artefacts. Pairs of clusters with similar waveforms and coordinated refractory periods in the cross-correlograms were merged into a single cluster. The odor response recording session and the opto-tagging recording session for each experiment were merged and sorted as one session and later split for subsequent analysis.

### Opto-tagging

Opto-tagging was performed at the end of each experiment (or at the start for experiments using DREADDs). When the probe without the attached optic fiber was used, a separate 200μm or 400μm optic fiber (ThorLabs) was implanted at a 12°angle to the probe, and a distance of 500-600μm away from the probe’s ventral coordinate. The optic fiber was connected to a mechanical shutter (Uniblitz, Vincent Associates, Rochester, NY) and then to either a 532nm or 635nm laser (OptoEngine LLC, Midvale, UT) using a patch cable (ThorLabs). The mechanical shutter was controlled using TTL pulses sent from an analog output of the Cerebus system which then allowed light pulses through the optic fiber (1000 pulses, 300ms, 1Hz). The same TTL pulse was also sent to an analog input channel on the Cerebus system to align light pulses with electrophysiological recordings. Latency to suppression was determined by change point analysis (Wolff et al., 2014, Rowland et al., 2018) performed using Matlab’s ‘findchangepts’ function. A cell was categorized as light-responsive if (1) its trial-averaged baseline activity was significantly higher than that during the laser pulse (Wilcoxon rank-sum test, p<1×10^-7^), (2) the area under the ROC (auROC) curve was less than 0.5 and (3) the changepoint value was within 10ms of the laser on time. In 4/9 optogenetic suppression experiments, we delivered 60 1-second light pulses which were triggered by a TTL pulse to the laser (i.e. no shutter), and thus we were not able to accurately determine change points. For these experiments, light-responsive cells were determined using the rank-sum test and auROC criteria, followed by manual curation. Units that exhibited ‘fast’ suppression were categorized as light-responsive, while units that exhibited visibly slow or delayed suppression were categorized as non-light responsive.

### Odor stimuli

The odor stimulus set consisted of 2-hexanone (Sigma, 02473), octanal (Aldrich, O5608), isoamyl acetate (Tokyo Chemical Industries, A0033), ethyl butyrate (Aldrich, E15701), valeraldehyde (Aldrich, 110132), ethyl tiglate (Alfa Aesar, A12029), acetophenone (Fluka, 00790), γ-terpinene (Aldrich, 223190), ethyl acetate (Sigma-Aldrich, 34858) and methyl tiglate (Alfa Aesar, A11964).

### Odor delivery

Odors were delivered using a custom-built 16-valve olfactometer. House air was passed through a filter and then split three ways through three mass flow controllers (MFCs) (Aalborg, Orangeburg, NY). MFC1 flowed filtered air at a constant rate of 1L/min while airflow through MFC2 and MFC3 were varied to vary the concentration of odors delivered while maintaining a total airflow out of the olfactometer at 1L/min. Custom Arduino software was used to control odor solenoid opening and closing as well as provide a log for which odor was delivered. Monomolecular odorants were diluted to 1% v./v. in mineral oil. Normally, a 1L/min clean air stream from MFC1 was directed to the mouse’s nose. During a trial, air from either MFC1 or MFC2 was directed through one of the odor vials and the odorized air stream was directed to exhaust for an equilibration period of 6 seconds before rapid switching of a final valve, triggered on exhalation, re-directed odorized air to the nose and neutral air to exhaust. Odors or mineral oil blank control stimuli were presented for 1 second, and stimuli were presented every 10 seconds in random order.

### Spontaneous firing rate

Spontaneous firing rates were determined by counting the number of spikes in a 4-second inter-trial window starting 4 seconds after odor offset.

### Individual cell-odor responses

Individual trial-averaged cell-odor responses as a function of time were visualized by computing kernel density functions smoothed with a 10ms Gaussian kernel. The reliability of responses was determined using the auROC metric. auROC values were computed from the U-statistic of the rank-sum test comparing spike counts of each cell-odor pair during the first sniff after odor presentation to spike counts during the sniff preceding odor presentation. auROC values were then converted to response indices using the following formula: au-ROC*2-1. A value of −1 or 1 indicates a suppressed or activated odor response that was perfectly distinguishable from the pre-odor baseline response while a value of 0 indicates no difference between the odor and preodor baseline response.

### Sparseness

Lifetime sparseness describes the firing rate distribution of individual neurons to all of the odors presented and reflects the tuning width of neurons. A value of 1 indicates the cell responded to just one odor and a value of 0 indicates the cell’s firing rate was distributed uniformly across all odors. Population sparseness is a measure of the density of responses of a population of cells, where a value of 1 indicates the population response to a given odor was driven by just one cell and a value of 0 indicates uniform contribution of all neurons in the population to a given odor response. Lifetime and population sparseness values were calculated for each experiment individually:

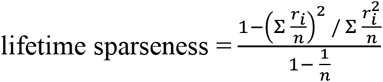

where r_i_ is the trial-averaged response to the *i*th odor and n is the number of odors. Population sparseness for each odor was calculated with the same formula but r_i_ is now the trial-averaged response of the *i*th cell and n is the number of cells.

### Trial-trial population vector distances and principal components analysis

The similarity of population odor responses, which are vectors of z-scored spike counts in the first 500 ms after inhalation onset, was determined by correlating response vectors from pairs of trials in each experiment separately. Spike counts were z-scored to a 500 ms epoch before inhalation onset (−.6 to −.1 s before inhalation onset (t=0)). The mean within-odor correlation was calculated by averaging correlation coefficients of trials from the same odor while the mean between-odor correlation was calculated by averaging correlation coefficients of all trials except those of the same odor. Principal component analysis was performed using Matlab’s built-in ‘pca’ function using z-scored pseudopopulation (responses from all experiments were pooled together) spike count vectors.

### Population decoding analysis

Odor classification accuracy was determined with a linear multiclass support vector machine (SVM) classifier using leave-one-out cross validation (LIBLINEAR, solver 4 – support vector classification by the Crammer and Singer method, https://www.csie.ntu.edu.tw/~cjlin/liblinear/). The feature vectors used in the classifier were pseudopopulation vectors of raw spike counts in the first 500 ms after inhalation onset. To determine classification accuracy at increasing numbers of cells, populations of varying sizes were constructed by subsampling the total pseudopopulation 100 times and averaging the classification accuracy across all 100 iterations for that population size.

### Latency analysis

Response latencies were determined for all active cell-odor pairs (response index > 0). For peak latencies, KDFs were first computed from spike counts occurring in a 300 ms time window from inhalation onset using a 10 ms Gaussian kernel. Then, the time of the maximum value of the KDF was identified and taken as the latency to peak. For onset latencies, KDFs were similarly computed but for a time window of 600 ms, spanning 300 ms before inhalation onset (baseline) and 300 ms after. The mean firing rate and standard deviation was determined for the baseline period and the time after inhalation onset at which the KDF first exceeded two standard deviations above the baseline was then taken as the onset latency. Histograms were constructed using 2 ms bins.

Odor classification accuracy in a sliding window was determined using a linear multiclass support vector machine (SVM) classifier using leave-one-out cross validation, as described in the ‘Population decoding analysis’ section. Within each time window, the pseudopopulation spike count vector for one trial was omitted and the mean across trials of the remaining responses for each odor was calculated. Classification within each window was repeated 100 times and accuracies for each window are the average across all iterations.

### DREADD agonists dosage and delivery

250 μL of either CNO (4 mg/kg, Sigma, C0832) or Compound 21 (1 mg/kg, Sigma, SML2392), diluted in sterile 0.9% saline solution was injected intraperitoneally during the recording session.

### Immunohistochemistry

Mice were anaesthetized with isoflurane and then perfused transcardially with cold 4% paraformaldehyde in 0.2 M phosphate buffer (4%PFA). Brains were dissected and post-fixed for 48 hours in 4% PFA at 4°C and then placed in a vibrating microtome, submerged in cold 1x phosphate buffered saline (PBS, Sigma, P4417) to slice 50μm sections of olfactory bulb and piriform cortex. To stain for GFP and mCherry, sections were permeabilized using 0.1% Triton X-100 (Acros Organics, 327371000) in 1xPBS (t-PBS) for 3 washes and then with 0.3% t-PBS for 1 wash. Slices were incubated with chicken anti-gfp (1:400, Invitrogen, A10262), rabbit anti-rfp (1:400, Rockland, 600-401-379) in blocking buffer (5% normal goat serum (EMD Millipore, S26) in 0.3% t-PBS overnight at 4°C. The next day, slices were rinsed in 0.1% t-PBS for 3 washes and then incubated in AlexaFluor 488 goat anti-chicken (1:400, Life Technologies, A11039), AlexaFluor 555 goat anti-rabbit (1:400, Invitrogen, A32732) and NeuroTrace 435/455 (1:300, Life Technologies, N21479) in blocking buffer overnight at 4°C. Slices were again washed in 0.1% t-PBS and then mounted and cover-slipped with Fluoromount-G. Flourescence images were taken using a Zeiss 780 inverted confocal microscope and adjusted for brightness and contrast using ImageJ.

## Author contributions

S.N. and K.M.F. conceived the experiments, analyzed the experiments, and wrote the paper. S.N. performed all experiments except slice electrophysiology experiments, which were performed by K.M.F.

## Acknowledgements

We thank K. Bolding for invaluable technical assistance and comments on the manuscript, L. Glickfeld, C. Hull, F. Wang and A. Sanzeni for valuable discussions, and R. Blazing, C. Diaz, A. Fleischmann, H. Jackson, F. Santos-Valencia, A. Schaefer and F. Wang for helpful comments on earlier versions of the manuscript. This work was supported by grants from the NIH (DC015525, DC016782, U19 NS112953), a Holland-Trice Scholar Award (K.M.F) and a Holland-Trice Graduate Fellowship (S.N.).

## SUPPLEMENTARY MATERIALS

**Supplementary Figure 1.**
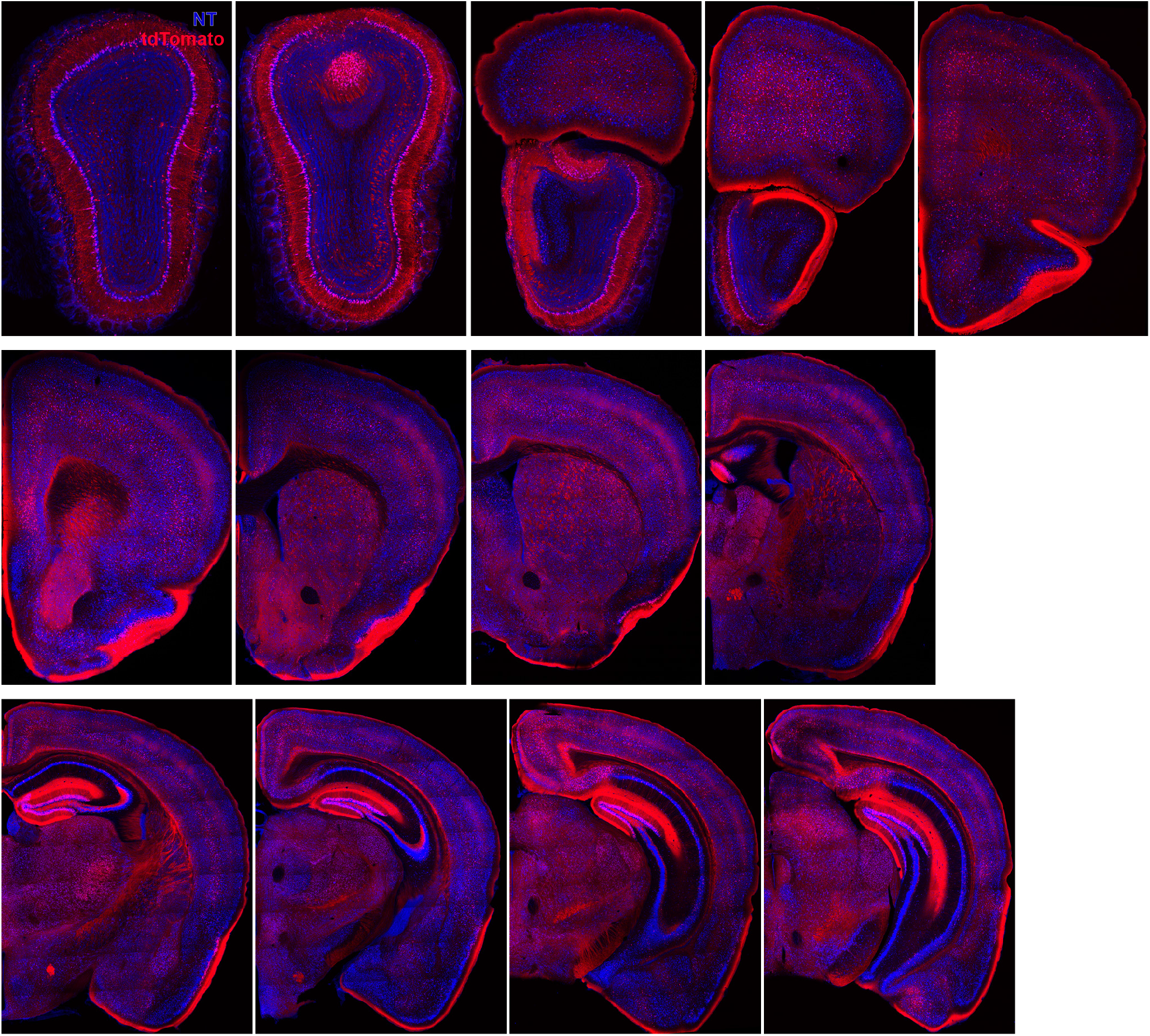
Ntng1+/Cre+ neurons throughout the brain. A series of coronal sections through a Ntng1-Cre/Ai14 mouse showing Ntng1 expression patterns.

**Supplementary Figure 2.**
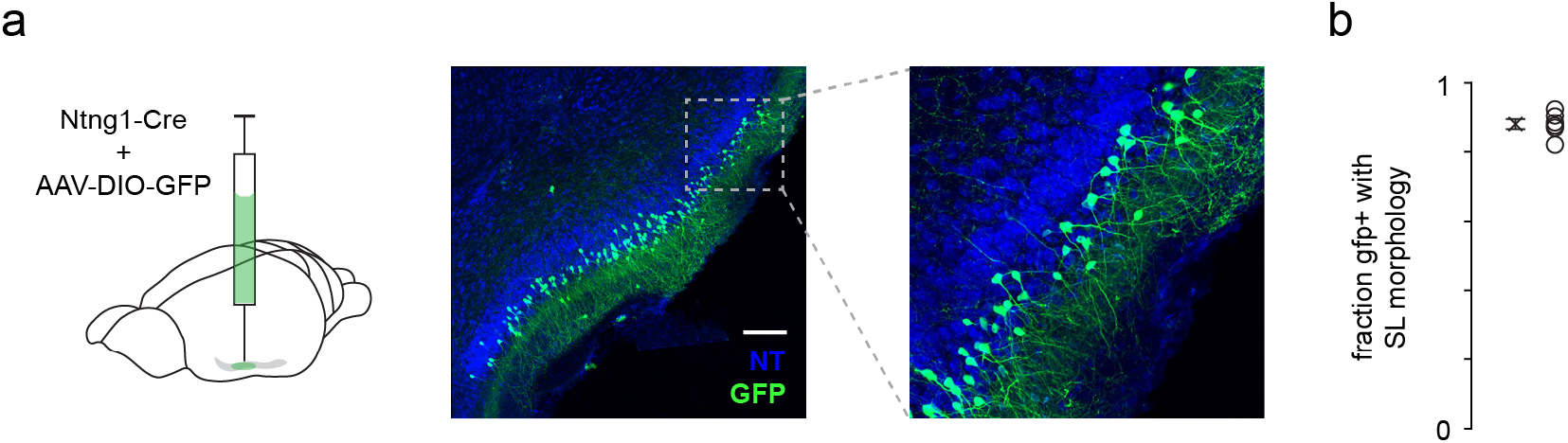
Ntng1+ cells have similar laminar and morphological properties as SLs. (**a**) Low titer of a Cre-dependent GFP was injected into anterior PCx to sparsely label a small subset of Ntng1+ neurons. This allowed better visibility or GFP+ cell morphology and laminar location. Scale bar is 200 μm. (**b**) Quantification of the fraction of GFP+ cells that have SL morphology (half-moon shaped somata, two prominent apical dendrites, no basal dendrites) and were located in Layer IIa.

**Supplementary Figure 3.**
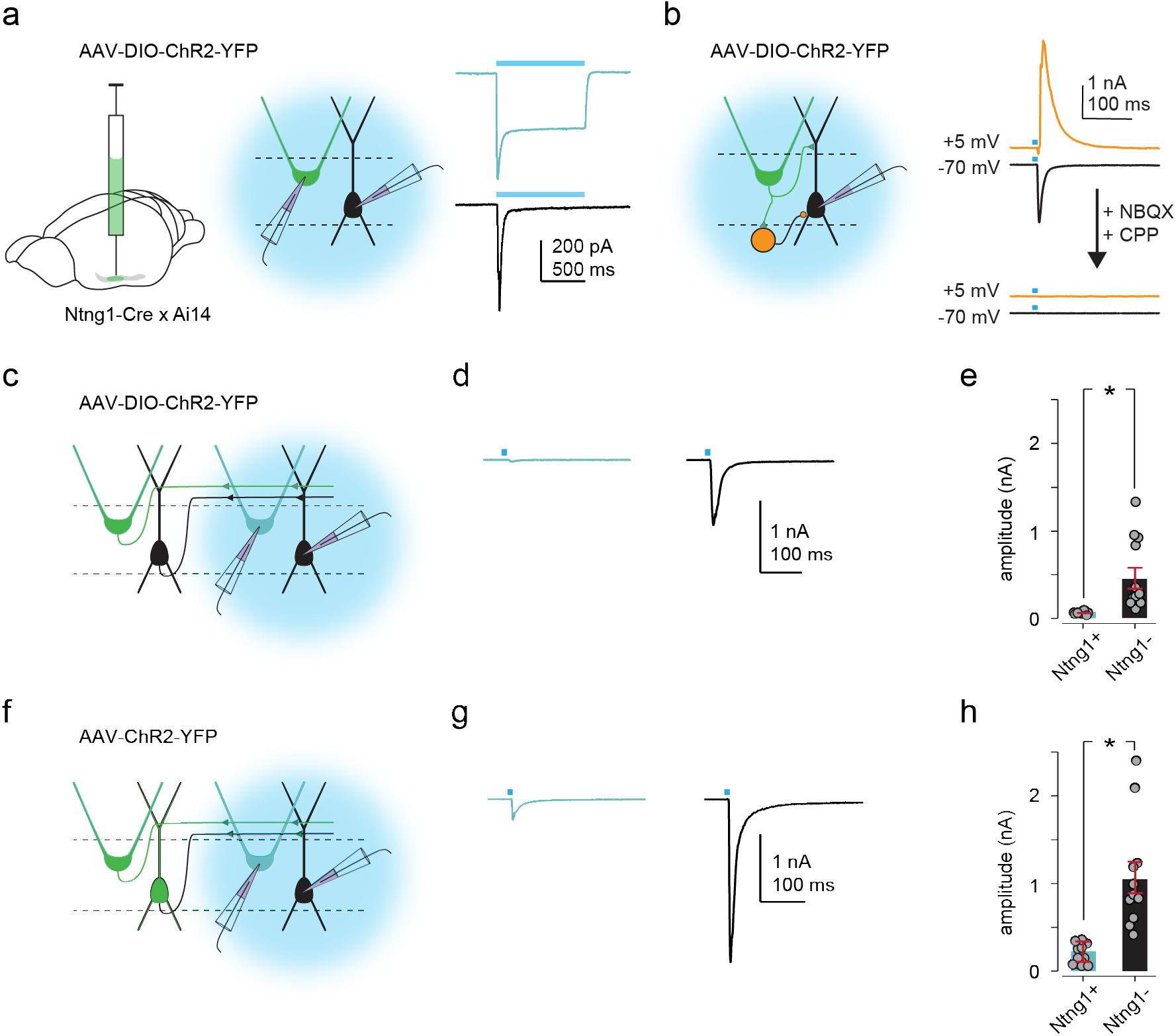
Ntng1+ and Ntng1- cells have similar intrinsic and synaptic connectivity properties as SLs. (**a**) Left and middle: experimental schematic. Cre-dependent ChR2 was injected focally into anterior PCx of Ntng1-Cre crossed with Ai14 mice. Whole-cell voltage clamp recordings were obtained from ChR2-expressing Ntng1+ (tdTomato+) and Ntng1- cells while illuminating the slice with blue (473nm) light. Right: large, sustained photocurrents were recorded in Ntng1+/tdTomato+ cells (blue) but not in Ntng1-/tdTomato-cells (black). Disynaptic EPSCs were recorded in Ntng1- cells. (**b**) Left: experimental schematic. Light-evoked synaptic responses in Ntng1- cells to strong, brief (1 ms) light pulses were recorded while holding the cell at −70 mV, to isolate excitatory postsynaptic currents, and at +5 mV, to isolate inhibitory postsynaptic currents. Right: light-driven activation of Ntng1+/ChR2+ cells evoked large, transient inward and outward responses in Ntng1- cells at −70 mV and +5 mV, respectively. Responses at both holding potentials were completely blocked by glutamate receptor antagonists (10 μM NBQX and 10 μM CPP). Ntng1+ cells are therefore glutamatergic. (**c**) Experimental schematic. As in (**a**) but whole-cell recordings were obtained from Ntng1+ cells that were located away from the injection site (i.e. cells that were not expressing ChR2). (**d**) Example EPSC traces evoked in Ntng1+/ChR2- (blue) and Ntng1- (black) cells by light-driven activation of Ntng1+/ChR2+ cells. (**e**) Summary plot of EPSC amplitude in Ntng1+ (25.3 ± 5.76 pA; n = 12 cells, 3 mice) and Ntng1- cells (429 ± 114 pA, n = 13 cells, 3 mice) in response to light-driven activation of Ntng1+/ChR2+ cells. (**f**) Experimental schematic. As in (**a**) and (**c**) but a non-conditional ChR2 was injected focally into anterior PCx. (**g**) Example EPSC traces evoked in Ntng1+/ChR2- (blue) and Ntng1-/ChR2- (black) cells by light-driven activation of all ChR2+ cells. (**h**) Summary plot of EPSC amplitude in Ntng1+ (187 ± 30.6 pA, n=14 cells, 3 mice) and Ntng1- cells (1060 ± 175 pA, n=12 cells, 3 mice) in response to light-driven activation of all ChR2+ cells.

